# New insights on the Feeding Site and Salivation by *Lutzomyia longipalpis* (Diptera: Psychodidae) During Blood Ingestion on Host Skin

**DOI:** 10.64898/2026.02.23.707437

**Authors:** Kelsilandia Aguiar Martins, Mauricio Roberto Viana Sant’Anna, Yevhen F Suprunenko, Luccas Gabriel Ferreira Malta, Adalberto Alves Pereira Filho, Ricardo Nascimento Araujo, Nelder Figueiredo Gontijo, Marcos Horácio Pereira

## Abstract

Phlebotomine sand flies are major vectors of *Leishmania* parasites, yet the mechanisms underlying their blood-feeding behavior remain poorly understood. In *Lutzomyia longipalpis*, the primary vector of *Leishmania infantum* in the Americas, feeding occurs via telmophagy, a pool-feeding method which is known by involving dermal laceration, salivation, and the creation of a blood pool. While the biochemical effects of sand fly saliva on host hemostasis, inflammation, and immunity are well studied, the dynamics of mouthpart movements and saliva at the feeding site remain to be systematically explored. Using intravital microscopy, fluorescent saliva labelling and image analysis, we characterized the mechanical actions of mouthparts and the spatial-temporal patterns of salivation during feeding on mammalian skin. Our recordings indicate that the labrum and hypopharynx are the most prominent mouthparts during feeding and exhibit scissor-like movements during probing. At specific moments, these structures close forcefully, generating small blood splashes in multiple directions. Feeding occurred in two distinct phases: an initial probing phase, often distinguished by ineffective blood intake, and a subsequent engorgement phase that was initiated exclusively upon the activation of small dermal “feeder vessels.”Acridine Orange labelling showed abundant early salivation that penetrated progressively deeper into the dermis and remained detectable for over an hour, reflecting both the tissue damage and enzymatic effects. The analysis of images demonstrated the sequential salivation events, highlighting an initial high-frequency phase followed by a more gradual pattern during engorgement. These findings provide the first real-time, detailed view of the coordinated interactions between mouthpart mechanics, targeted salivation, and host microvascular responses in *Lu. longipalpis*. This study redefines sand fly telmophagy as a non-passive and coordinated process integrating mouthpart mechanics, salivation, and modulation of host vasculature. This work advances our understanding of sand fly vector-host interactions and underscores the potential of salivary molecules as targets for transmission-blocking strategies.

**Author Summary:** Phlebotomine sand flies are the main vectors of *Leishmania infantum*, the parasite responsible for visceral leishmaniasis in the Americas. Although sand flies are traditionally classified as “pool feeders,” meaning they lacerate the skin and feed from small pools of blood, the mechanics of how they obtain blood and deliver saliva into host skin have remained poorly understood. In this study, we used image analysis, intravital microscopy and fluorescent labeling of saliva to visualize, in real time, the feeding behavior of *Lutzomyia longipalpis* on mammalian skin. We show that blood feeding is not a passive process based solely on blood pooling. Instead, it involves coordinated movements of the mouthparts, modulation of host microvessels with the saliva contribution, and the recruitment of small dermal “feeder vessels” that supply blood directly to the insect. Our findings reveal that sand fly feeding is a highly orchestrated interaction between vector and host, integrating mechanical tissue disruption, salivary secretion, and vascular responses. These processes likely create a favorable microenvironment for *Leishmania* establishment and transmission. By providing a detailed characterization of mouthpart and salivation dynamics, this study advances our understanding of sand fly biology and highlights salivary components and feeding-site events as potential targets for transmission-blocking strategies.

## Introduction

Phlebotomine sand flies are small insects comprising over 800 species distributed worldwide (1). Approximately 10% of these species are involved in the transmission of pathogens to humans, including the ones that cause leishmaniasis, bartonellosis and pappataci fever. Among them, *Lutzomyia longipalpis* (2) stands out as the principal vector of *Leishmania infantum* (Kinetoplastida: Trypanosomatidae), the etiological agent of visceral leishmaniasis (VL) in the Americas (3). Recent estimates indicate that 600,000 to 1 million new cases of cutaneous leishmaniasis occur globally each year, alongside 50,000 to 90,000 new cases of visceral leishmaniasis (4). Approximately 90% of visceral leishmaniasis cases are concentrated in the Indian subcontinent, East Africa, and Brazil (5). Like other vector-borne diseases, leishmaniasis involves the transmission of parasites from invertebrates to vertebrates during the blood-feeding process, which typically occurs within the dermis of the vertebrate host (6). In their adult stage, both male and female sand flies feed on plant juices or sugary secretions to meet their metabolic energy demands. However, blood feeding is an activity performed exclusively by adult females and is essential for reproductive success, as it provides proteins and amino acids required for oogenesis (7),(8).

The sand fly *Lu. longipalpis* possesses rigid mouthparts. Its proboscis is typically short (<1 mm) and composed of the labrum, hypopharynx, paired mandibles, and laciniae (9). Similarly to tabanids, ceratopogonids, tsetse flies, and blackflies, sand flies are recognized for a blood-feeding strategy known as telmophagy or “pool feeding” (10), (11). This method involves inserting the mouthparts into the host’s skin, rupturing superficial blood vessels in the dermis, and creating a small pool of blood. Blood is then ingested through the action of cibarial and pharyngeal pumps and transported to the midgut, where blood digestion occurs (12), (13).This feeding strategy is considered highly effective for *Leishmania* transmission, as the parasites primarily reside within skin macrophages rather than circulating freely in the bloodstream (14).

Regardless of the feeding mechanism - solenophagy (vessel feeding) or telmophagy (pool feeding) - the movement of arthropod mouthparts while probing for blood damages the skin and endothelial lining of blood vessels, thereby triggering a cascade of host responses. These include platelet aggregation, vasoconstriction, coagulation, increased vascular permeability, and leukocyte chemotaxis. Additionally, hemostatic and inflammatory reactions may be exacerbated by the host’s immune responses (both innate and adaptive) against salivary antigens released during the blood meal (15),(16).

To counteract these host defenses, hematophagous arthropods inject saliva containing a variety of bioactive molecules that modulate hemostasis, inflammation, and immunity, thereby facilitating blood acquisition (17),(18). Fleas and most blood-feeding nematocerans (including mosquitoes, sand flies, and blackflies) are estimated to possess between 100 and 200 salivary proteins, whereas brachycerans (e.g., tsetse flies and tabanids) have approximately 250 - 300, triatomines over 300, and ticks more than 500 (17), (19).These differences in salivary complexity likely reflect variations in feeding strategies (vessel vs. pool feeding) and the duration of the blood meal (19).

Beyond aiding blood feeding, the saliva of disease vectors, as demonstrated in several experimental models (20) (21),(22),(23), (24),(25) can also facilitate pathogen transmission. The first evidence supporting the role of sand fly saliva in *Leishmania* infectivity emerged in the early 1980s. Since then, numerous salivary components, particularly those with immunomodulatory functions, have been characterized. Importantly, some of these components hold promise for inclusion in future leishmaniasis transmission blocking vaccine formulations (21), (26).

Recent findings indicate that alterations at the feeding site arising from mechanical injury caused by mouthpart movements and the deposition of saliva play a pivotal role in shaping *Leishmania* infection dynamics. These observations highlight the feeding site itself as a potential niche for parasites, thereby facilitating their successful establishment and subsequent transmission to vertebrate hosts (27),(28), (26).

Although the mechanical aspects of blood acquisition and the role of saliva in Leishmania transmission are well established, a substantial knowledge gap persists regarding the dynamics of blood acquisition and salivation in sand flies during their feeding on host skin. This gap motivates the present study, which aims to characterize the mechanical properties of blood feeding and the salivation patterns of *Lu. longipalpis* on mammalian skin.

## Materials and Methods

### Insects

A colony of *Lu. longipalpis* sand flies, originally established from specimens collected in Teresina, Brazil, was maintained according to the protocol described by (29). The colony was reared in the insectary of the Laboratory of Physiology of Hematophagous Insects, Department of Parasitology, ICB/UFMG, under controlled environmental conditions (Temperature: 25°C, under 70% ± 10% relative humidity, and 12 h light / 12 h dark photoperiod). All experiments were conducted using groups of up to five female sand flies aged between 4-8 days post-emergence.

### Mammalian Host

Four-to-five-week-old hairless mice (HRS/J strain) were obtained from the Animal Care Facilities of the Department of Parasitology, ICB/UFMG. The animals were housed in standard laboratory cages, maintained under controlled conditions, and provided food and water ad libitum. All mice used in the experiments had no previous exposure to sand flies. Experimental procedures were conducted in accordance with institutional guidelines and were approved by the Ethics Committee for Animal Use of the Federal University of Minas Gerais (CETEA/UFMG; protocol number 316/2015).

### Saliva Labeling

To fluorescently label sand fly saliva, newly emerged females were allowed to feed *ad libitum* on cotton pads soaked in a 30% sucrose solution containing 0.1% Acridine Orange (AO). Salivary glands were dissected 48 hours after exposure and examined under an epifluorescence microscope equipped with a filter set for AO fluorescence (excitation: 450–480 nm; emission: 520 nm) to confirm dye uptake by the glands.

### Measurement of Salivary Gland pH

To assess the pH of salivary glands, five females aged 3 to 6 days post-emergence were dissected in Insect Saline (IS: 119.7 mmol·L⁻¹ NaCl, 2.68 mmol·L⁻¹ KCl, 1.36 mmol·L⁻¹ CaCl₂, and 0.56 mmol·L⁻¹ glucose). Each *L. longipalpis* salivary gland is composed of two acini. One acinus per insect was transferred to a microplate well containing a pH indicator solution. Two pH indicators were used: bromothymol blue (Sigma-Aldrich, code 18470; pKa = 7.4) and bromocresol purple (Sigma-Aldrich, code 860891; pKa = 6.3). Both dyes were prepared as saturated solutions in IS (30). The pH of the bromothymol blue solution was adjusted to ≥7.0, and that of the bromocresol purple solution to ≤5.0 (31), (32). Following transfer of the acinus into the indicator solution, glands were mechanically ruptured, and color changes in the surrounding medium were observed and documented using photomicrography.

### Intravital Microscopy and Feeding Chamber

Intravital microscopy was performed using the method described by (33), with adaptations to accommodate sand fly feeding. An acrylic feeding chamber (5 cm diameter) was fabricated with a central 5 mm diameter aperture on the top surface. The mouse ear was positioned over this opening and secured using double-sided adhesive tape, allowing the inner ear dermis to be accessible to the insects. A second lateral port was included to facilitate the controlled release and removal of the insects (Figure 1). Mice were anesthetized via intraperitoneal injection of ketamine (150 mg/kg; Cristália, Brazil) and xylazine (10 mg/kg; Bayer, Germany). Core body temperature was maintained by keeping mice on a heating pad at 34 °C (Fine Science Tools Inc., Canada). After positioning the dorsal surface of the ear on the chamber aperture, a 30-minute equilibration period was observed to allow thermal and microcirculatory stabilization.

**Figure 1.**
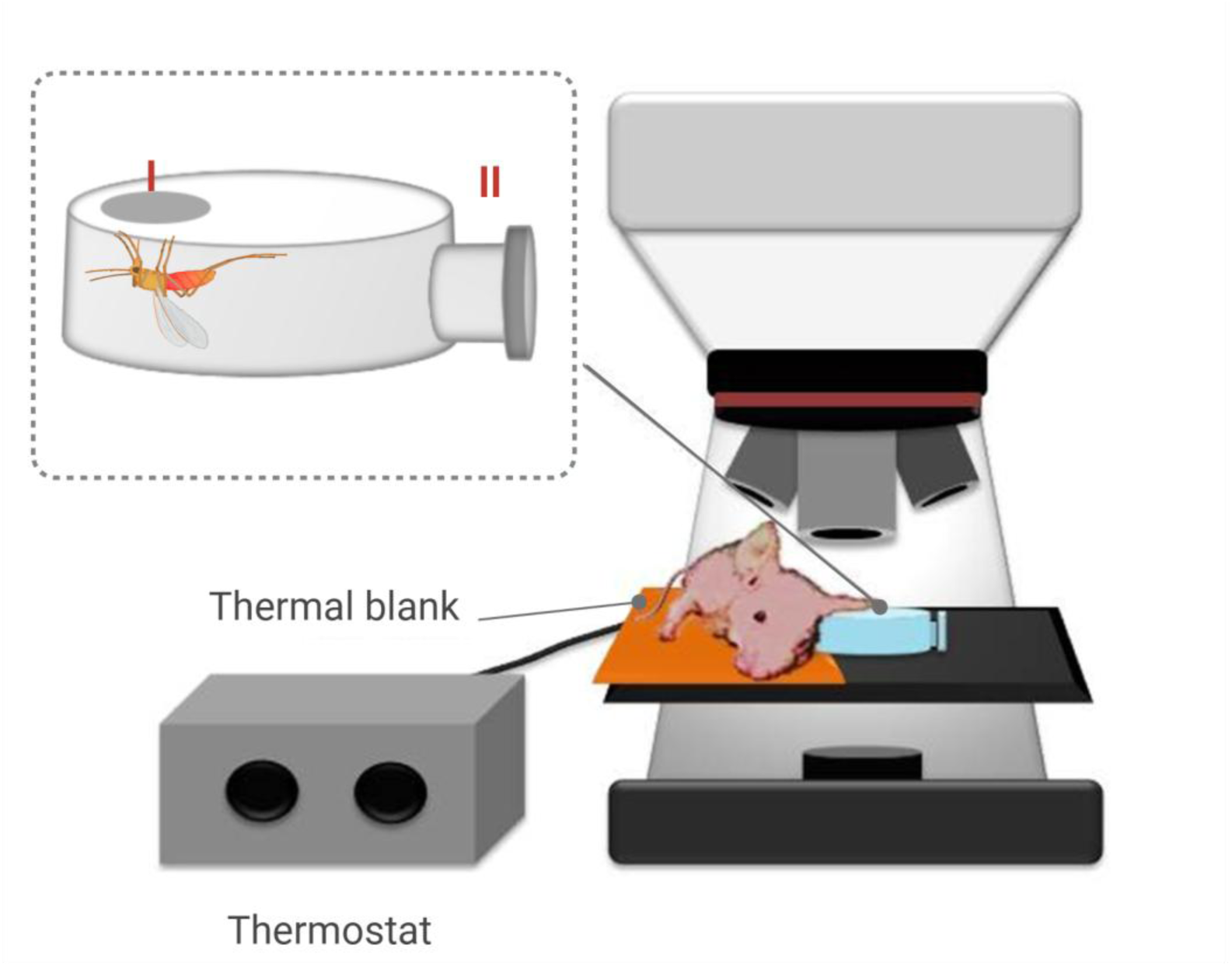
Schematic representation of the acrylic feeding chamber used for intravital microscopy of blood feeding by *Lutzomyia longipalpis* on the mouse ear. The chamber configuration includes an opening for ear exposure (I) and a lateral access port for insect manipulation (II).

Insect feeding was observed using an epifluorescence microscope (Leica DM500, Germany) equipped with a digital camera (Canon EOS 600D, Japan) to capture images and videos at 59.94 fps. In certain experiments, white-light transillumination was used to improve visualization of dermal microcirculation during sand fly probing and feeding.

### Image Analysis

Videos were analyzed using Fiji/ImageJ software (version 1.54f) (34). Fluorescence image processing followed the methods described by (35).To assess the spatial and temporal distribution of saliva release, the “Grouped Z Project” tool in Fiji/ImageJ was employed. This tool enables the division of image stacks into sequential groups of slices, each of which is processed independently using projection algorithms. Two projection types were applied: “Sum Slices” to highlight cumulative fluorescence intensity, and “Standard Deviation” to emphasize temporal variation within slice groups. The resulting projected stack allowed dynamic visualization of salivary secretion events over time. Graphical representations of the processed data were generated using Microsoft Office Excel 365.

### Statistical Analysis

All statistical analyses were performed using GraphPad Prism (GraphPad Software, San Diego, CA, USA). Data normality was assessed using the Kolmogorov–Smirnov test. For datasets that met the assumptions of normal distribution, comparisons between groups were conducted using the unpaired Student’s t-test. When normality assumptions were not satisfied, non-parametric analyses were performed using the Mann–Whitney U test. Differences were considered statistically significant at P < 0.05.

## Results

### Sand Fly Feeding Site: Alterations in Host Skin Microvasculature

Upon landing on the host skin, some *Lutzomyia longipalpis* females initiated biting behavior within a few seconds. Among the mouthparts, the labrum and hypopharynx, being the most prominent and structurally distinct elements, were the most visible during the probing and engorgement phases of feeding (Figs 2B and G). Following skin penetration, these structures performed coordinated penetration and withdrawal movements, remaining closely apposed but occasionally separating in characteristic “scissor-like” motions (Figs 2A and B). During these events, small hemorrhagic spots became visible on the skin surface, some of which were associated with active blood uptake. Females appeared to attempt blood acquisition from these hemorrhagic pools; however, passive blood leakage (even in cases of significant bleeding) did not enable a prolonged feeding, instead resulting in erratic, short-lived sucking attempts.

**Figure 2.**
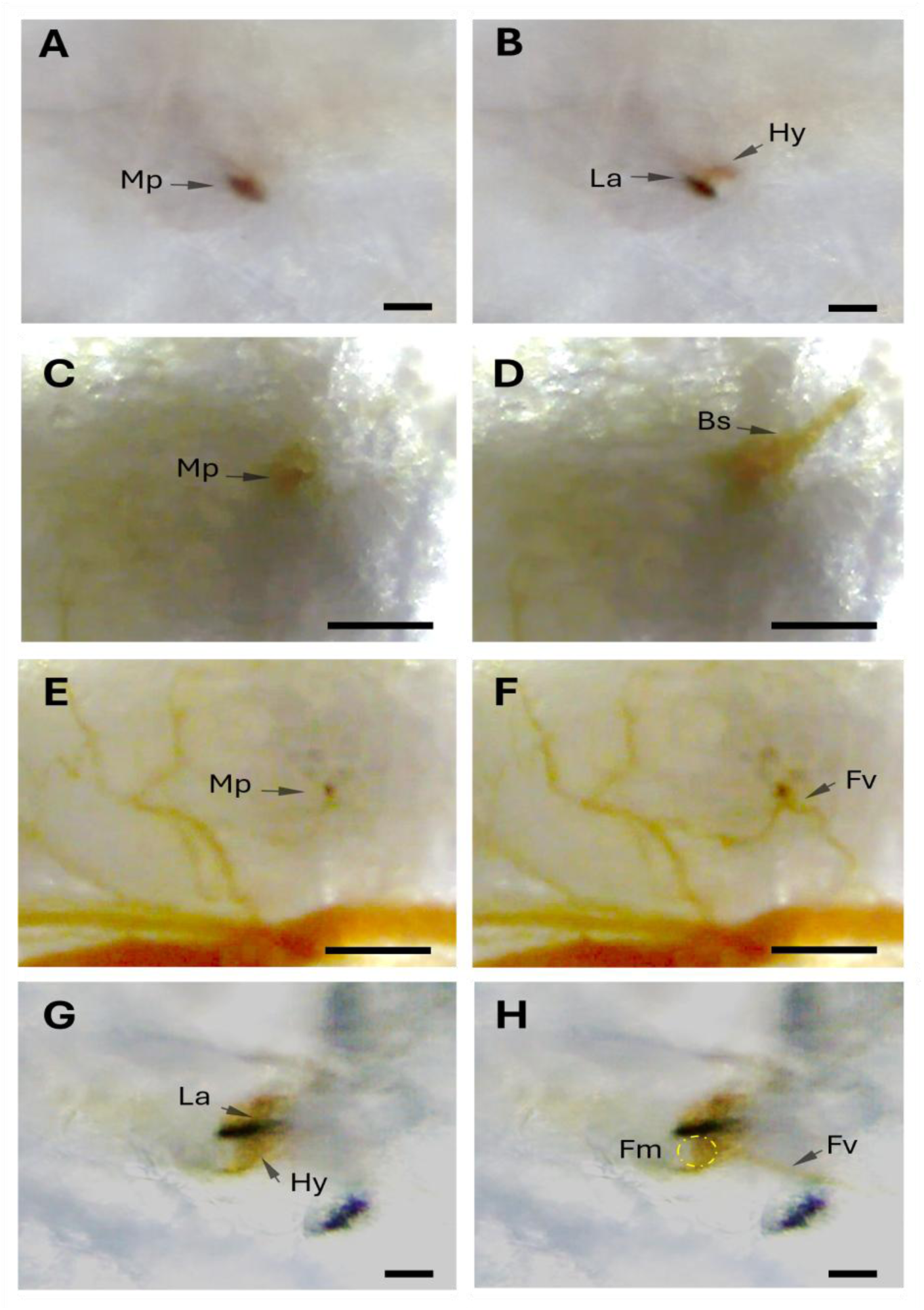
Mouthpart movements during probing and blood ingestion by *Lutzomyia longipalpis* in the mouse ear. (A–B) Following skin penetration, the mouthparts perform repeated penetration and withdrawal movements while remaining closely apposed (A) or transiently separating in characteristic scissor-like motions (B). (C–D) Emission of blood splashes from the mouthpart region during probing. (E–F) Vasodilation and opening of a “feeding vessel” supplying blood to the mouthpart region (red arrow), shown before (E) and after dilation (F). (G–H) Characteristic positioning of the labrum and hypopharynx during blood ingestion, illustrating moments of mouthpart closure (G) and association with the opened feeding vessel (H). Abbreviations: Blood splashes (Bs); Feeding vessel (Fv); Functional mouth (Fm); Labrum (La); Hypopharynx (Hy); Mouthparts (Mo); Bars=100µM.

Interestingly, in several instances during probing, the labrum and hypopharynx were observed to close forcefully, producing small blood splashes that dispersed in multiple directions (Figs 2C and D; S1 Video). Both the “scissor-like” movements and the blood splashes described above were frequently observed during transient interruptions in blood ingestion or during shifts in feeding sites, without withdrawal of the mouthparts from the host skin.

The process of blood feeding by *Lu. longipalpis* also induced a distinct sequence of alterations in the microcirculation of the host’s skin. Initially, insertion and mechanical movement of the mouthparts caused a transient vasoconstriction response (Figs 3A and B), during which small blood vessels became temporarily invisible under microscopic observation. This was followed by gradual re-establishment of blood flow in one or more vessels (Figs 2G and H), typically preceding the onset of sustained blood suction and engorgement by the insect. At a certain point, individual females abruptly ceased probing and entered a prolonged engorgement phase, during which they typically remained until completing the blood meal. As blood feeding continued, the most common observation was vasodilatation in the microcirculation at the feeding site (Figs 3C and D). During this phase, a directed blood flow was observed emanating from one or more small dermal vessels - here termed as “feeding vessels”- towards the insect’s mouthparts. In several instances, the initiation of engorgement was seen to coincide precisely with the “opening” of a feeding vessel (Fig s 2E and F). Notably, these vessels appeared to be ruptured, yet they did not collapse or form a pool, maintaining an intermittent flow pattern (S2Video), contributing to a stable and localized source of blood for ingestion. Only towards the end of the blood meal, interruptions in blood intake became more pronounced, accompanied by a progressive accumulation of blood around the feeding site. In many cases, these events culminated in a local hemorrhage, which occurred predominantly following the withdrawal of the mouthparts from the host skin (Fig 3F).

**Figure 3.**
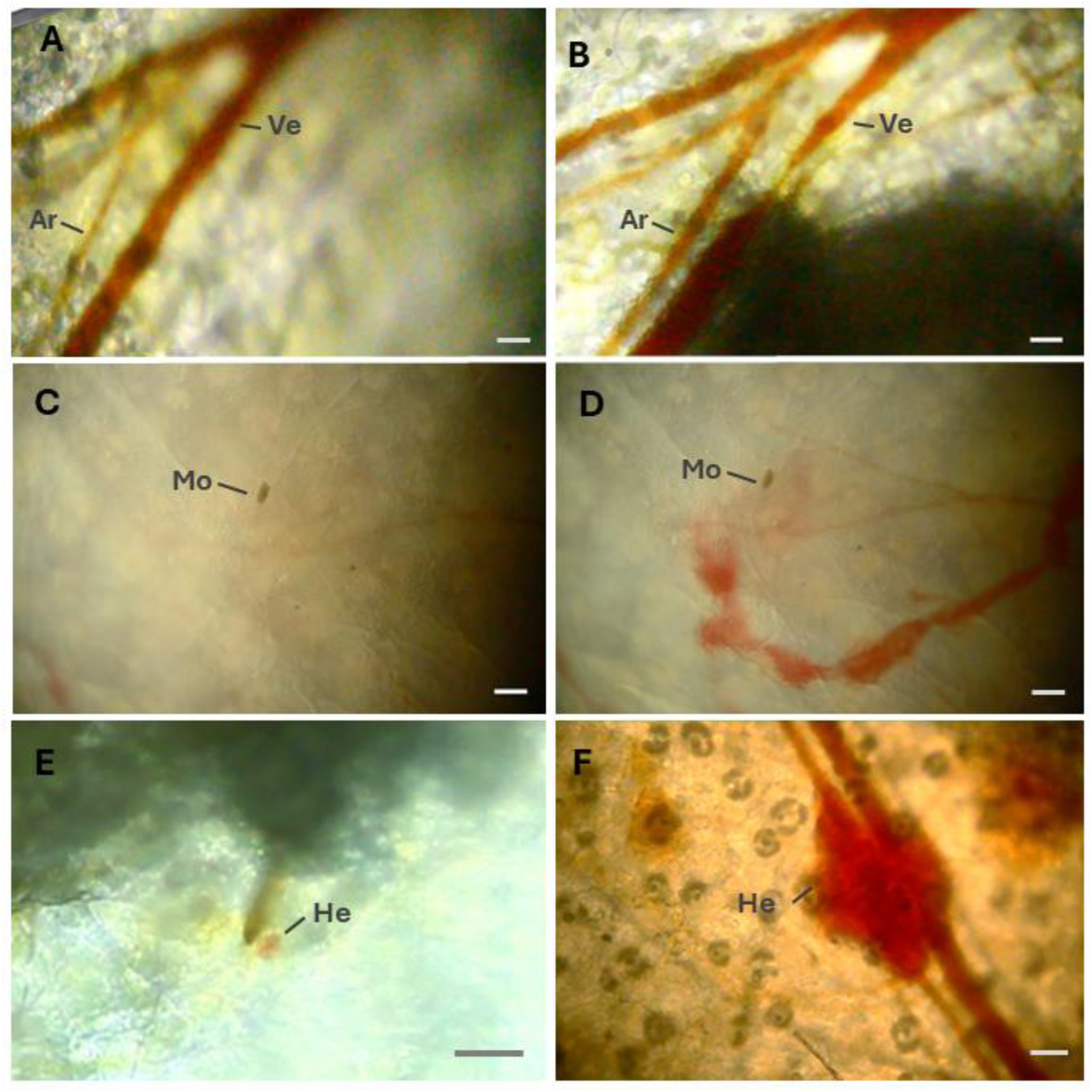
Vascular alterations at feeding sites during blood feeding by *Lutzomyia longipalpis.* (A) Microcirculatory architecture prior to the onset of feeding. (B) Typical vasoconstriction observed in vessels adjacent to the site of mouthpart insertion. (C) Early feeding phase, within the first seconds after probing initiation. (D) Pronounced vasodilation occurring during active blood ingestion. (E) Punctate hemorrhage commonly observed during the initial probing phase. (F) Extensive hemorrhage typically associated with mouthpart withdrawal at the end of the blood meal. Abbreviations: Arteriole (Ar); Hemorrhage (He); Mouthparts (Mo); Venule (Ve); Vasoconstriction (Vc); Bars=100µM.

### Saliva Labeling and Salivary Gland pH Measurement

Feeding female sand flies with a 30% sucrose solution containing 0.1% Acridine Orange (AO) resulted in strong and efficient fluorescence labeling of both the diverticulum and the contents of the salivary glands. Fluorescence was consistently observed within the salivary glands of females before and after the blood feeding (Figs 4A and B; Figs 4C and D) in all insects examined 48 hours post-exposure (n = 20). Effective labeling of the salivary glands was also achieved in insects that had previously been fed with sucrose alone prior to AO exposure (n = 10) and the intensity of labeling in the salivary glands was comparable regardless of whether AO was administered during the first or subsequent sucrose feedings. The survival rate of sand flies up to 10 days after ingestion of the AO-supplemented sucrose solution (n = 30) did not differ significantly from that of control insects fed on sucrose without AO (Mann-Whitney Test, n.s.).

**Figure 4-.**
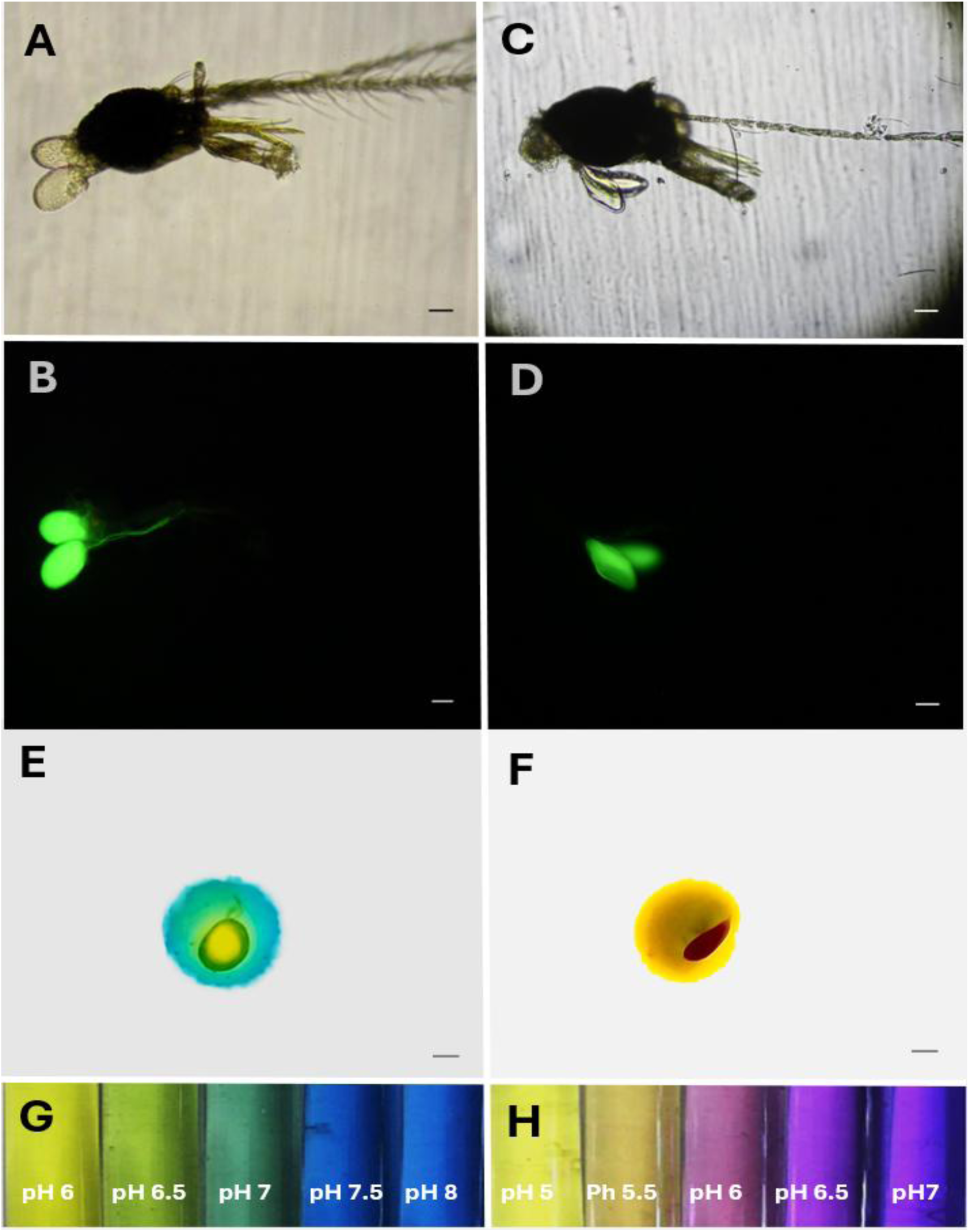
Salivary glands of *Lutzomyia longipalpis* stained with acridine orange and exposed to pH indicator solutions. (A–B) Paired salivary glands from a starved female visualized under light microscopy (A) and ultraviolet fluorescence (B) after staining with 0.1% acridine orange. (C–D) Salivary gland acini observed under light microscopy (C) and ultraviolet fluorescence (D), illustrating depletion of salivary contents following blood feeding. (E) Salivary gland acinus immersed in bromothymol blue, indicating an intraluminal pH ≤ 6.0. (F) Salivary gland acinus immersed in bromocresol purple, indicating a pH range of 5.5–6.0. (G) Buffered standard solutions with a pH range of 6 to 8 incorporating 0.1% Bromothymol Blue (H) Buffered standard solutions with a pH range of 5 to 7 incorporating 0.1% Bromocresol Purple; Bar scale=100µM

The pH of the salivary gland contents was estimated by immersing isolated acini of *Lu. longipalpis* in solutions containing pH indicator dyes—bromothymol blue and bromocresol purple. The observed color change of the indicators suggested that the pH of the salivary gland content was approximately 6.0 (Figs 4E and F; Figs 4G and H).

### Salivation by *Lu. longipalpis* in the Host Skin

The successful fluorescent labeling of sand fly saliva enabled clear visualization of salivary release in the host skin, beginning immediately after each bite and continuing throughout all phases of blood feeding. Saliva was observed exiting the salivary canal near the tip of the mouthparts - presumably the hypopharynx - and during penetration of the superficial layers of the ear skin. In these specific instances, a rapid transformation in saliva consistency was noted; the labeled (and likely unlabeled) saliva acquired a gel-like appearance immediately upon exposure to air.

Fluorescence analysis demonstrated that salivation persisted throughout the entire blood-feeding process. Monitoring fluorescence at the feeding site following insect detachment (n = 10) revealed that the signal remained detectable for a prolonged period (97.4 ± 12.7 min). In most assays, the intensity and persistence of fluorescence at the bite site hindered or completely obscured the direct visualization of rhythmic salivation, particularly during the early phases of feeding. In some experiments, it was possible to observe the labelled saliva released after the sandfly *Lu.longipalpis* punctures the host’s skin

To overcome the limitation caused by the fluorescence at the feeding site, two Fiji/ImageJ post-processing tools were applied to image sequences: (i) the Standard Deviation projection implemented through the Z Project function, generating a composite image representing the pixel-wise standard deviation across slices; and (ii) frame-by-frame subtraction (current frame minus the preceding frame) conducted using the Image Calculator tool. Figure 5A presents a comparative view between original 8-bit images (above) of the feeding site and the processed output obtained via frame subtraction (below). These analytical procedures enabled temporal segmentation and projection of image stacks, allowing more accurate assessment of the rate and spatial dynamics of saliva release throughout the different phases of the blood-feeding process.

**Figure 5.**
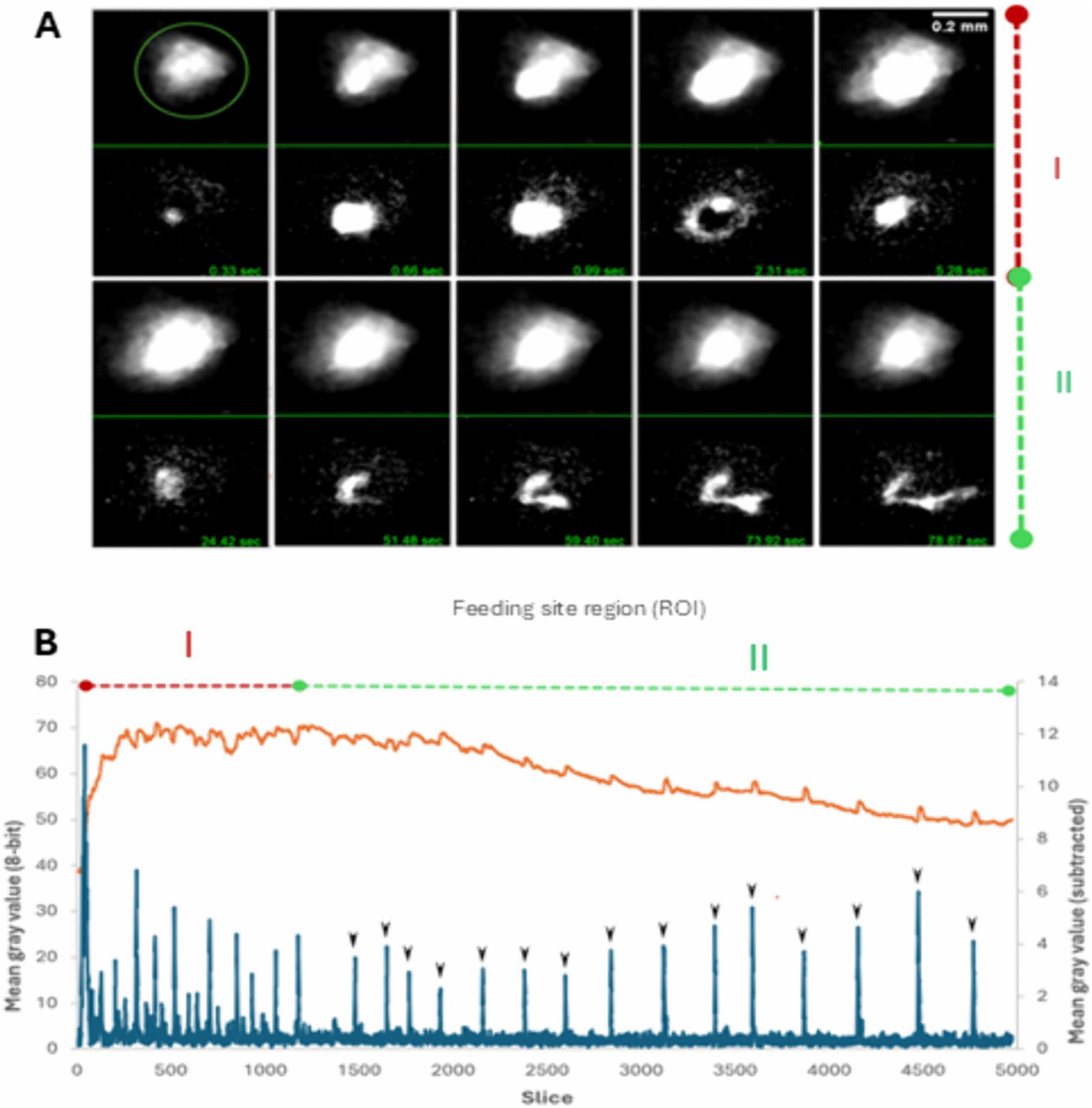
Saliva deposition by *Lutzomyia longipalpis* labeled with the fluorochrome acridine orange (0.1%) at the feeding site in mammalian skin. (A) Spatiotemporal visualization of saliva release at the feeding site over the course of blood feeding, represented here in 20 sequential time points. Upper panels correspond to original 8-bit images, whereas lower panels show processed images generated by frame subtraction using the Image Calculator tool (current frame minus the preceding frame). (B) Temporal variation in mean gray value measured at the feeding site region of interest (ROI; outlined in green in the first upper-left image) during sand fly hematophagy. Quantifications were obtained either directly from the original 8-bit image series (orange trace) or from frame-subtracted images produced via the Image Calculator tool (blue trace). Legend: Phase I (red) denotes the initial feeding phase, characterized by higher rates of saliva release. Phase II (green) corresponds to a subsequent phase in which salivary peaks occur more rhythmically and at lower frequency. Black arrows indicate discrete peaks of saliva deposition.

Using these image-processing approaches, two distinct patterns of salivation during sand fly blood feeding were identified. (a) Immediately following skin penetration, a substantial discharge of saliva was observed (Fig 5A, Phase I). This initial release was followed by a probing phase characterized by frequent peaks of salivary deposition (Fig 5B, Phase I), often concomitant with pronounced mouthpart movements. These movements were likely associated with repositioning of the mouthparts during the establishment of the functional feeding site—i.e., the tunnel-like structure formed within the host skin. (b) This initial phase was succeeded by a typically longer period extending until completion of the blood meal, during which salivary release occurred more rhythmically and at lower frequency (Fig 5B, Phase II). During this latter phase, the volume of saliva released per peak was reduced and tended to disperse from the primary site of deposition (Fig 5A, Phase II).

Complete monitoring of the salivation process was not feasible in all experiments. In females exhibiting sustained salivation for periods exceeding 1 min, fluorescence peaks displayed a mean frequency of 0.28 ± 0.13 Hz (n = 7). In several observations, however, insects did not remain stationary at a single feeding locus. Instead, they frequently initiated probing at one site and subsequently altered the axis of mouthpart movement, thereby expanding the area of saliva deposition. In other instances, females withdrew their mouthparts and either departed or resumed probing at a different location on the host skin. As previously described, the sudden closure of the mouthparts was also associated with the expulsion of saliva in jet-like formations toward the periphery of the feeding site. Image analysis from an experiment involving an insect with fluorescently labeled saliva (S3Video) revealed that these jets were typically preceded by the appearance of a fluorescence halo surrounding the mouthparts (Fig 6B) specifically the labrum and hypopharynx - followed by a rapid closing movement (Fig 6A). It was not uncommon to observe these saliva jets oscillating back and forth from the mouthpart’s region (S3Video). Over the course of the feeding event, the trajectory of these jets tended to lengthen, with some eventually continuing outward without returning toward the mouthparts (S4 Video S4).This pattern may be linked to the enzymatic action of hyaluronidase present in the insect’s saliva, which facilitates the cleavage of hyaluronic acid in the host dermal extracellular matrix, promoting deeper saliva diffusion within the skin.

**Figure 6.**
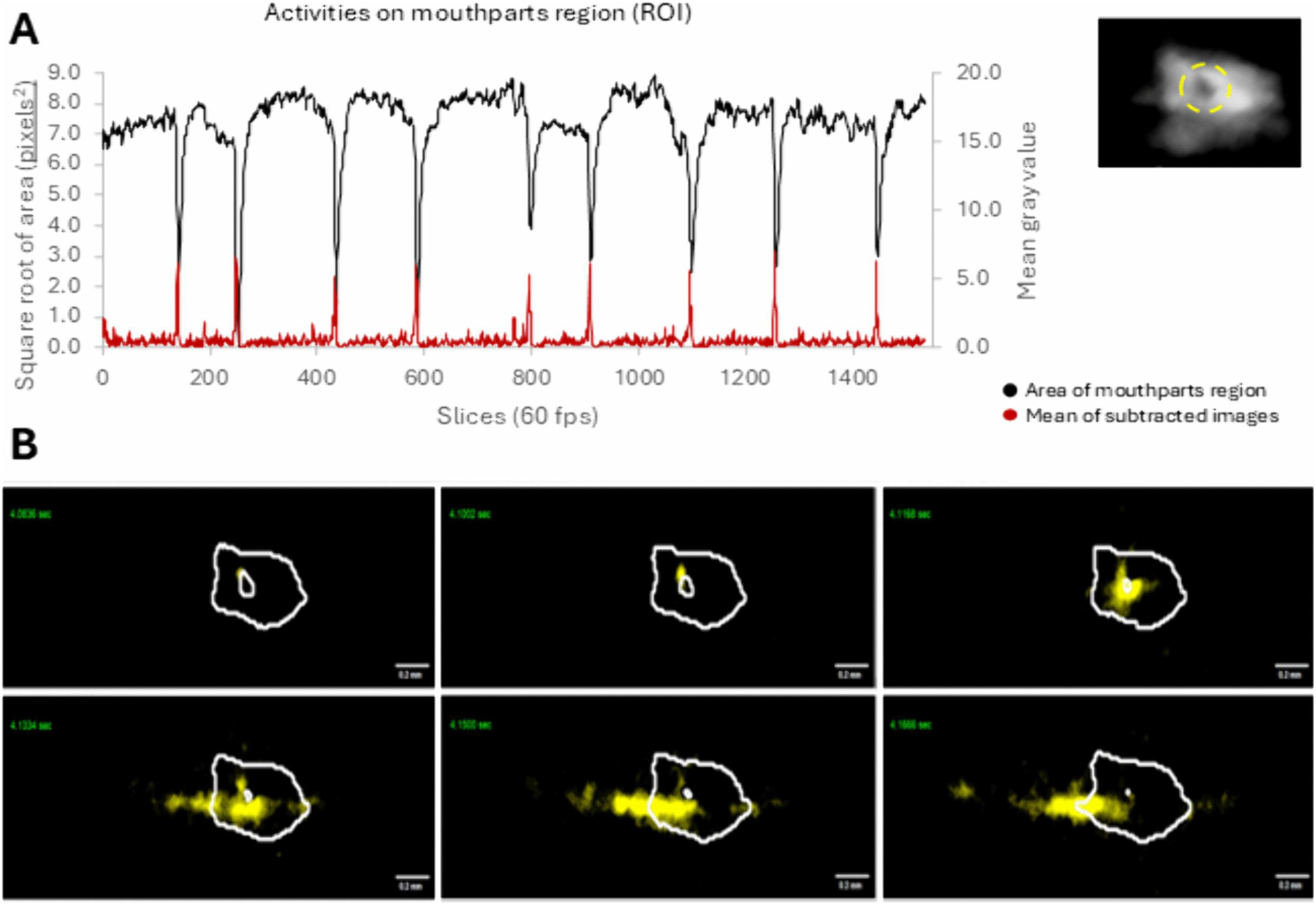
Sequential resolution of salivation events by *Lutzomyia longipalpis* through post-processed image sequence analysis. (A) Relationship between saliva emission (red-orange trace) and mouthpart region area (blue–black trace) in *Lutzomyia longipalpis* labeled with the fluorochrome acridine orange (0.1%) during blood feeding on host skin. Both parameters—saliva emission and mouthpart movement—were quantified within a region of interest (ROI) positioned adjacent to the mouthparts and highlighted in yellow in the representative image (upper right inset). Mouthpart area was measured from original 8-bit images using the Threshold and Analyze Particles functions, whereas saliva emission was quantified from the same image stack via frame-to-frame subtraction (current frame minus the preceding frame) performed with the Image Calculator tool. (B) Spatiotemporal representation of mouthpart movements and saliva deposition (yellow), including subsequent spreading, across six consecutive time points (0.0166 s intervals) during feeding. Masks corresponding to the mouthpart-occupied estimate area were generated from 8-bit images using the Threshold and Find Edges functions. Composite images were produced by combining frame-subtracted sequences with mouthpart masks through the OR operation using the Create New Window option in the Image Calculator tool. Red arrows indicate the mouthpart region, white arrows denote saliva deposition sites, and the white horizontal double-headed arrow represents saliva spreading away from the mouthpart region.

Application of the Grouped Z Project function for post-processing of image sequences obtained from *Lutzomyia longipalpis* females labeled with acridine orange (AO) revealed that salivary jets could penetrate ruptured, yet non-collapsed, microvessels located adjacent to the feeding site—structures consistent with so-called “vessel feeders” (Fig 7A–E). Interestingly, in some experiments in which females fed in close proximity to the walls of larger dermal vessels, fluorescent salivary jets were observed entering the vascular lumen and subsequently being transported by the bloodstream (Figs 7F–L; S5Video). Notably, in these cases, saliva appeared to traverse the vessel wall without overt structural rupture or detectable hemorrhage, suggesting the possibility of transvascular passage of salivary components.

**Figure 7.**
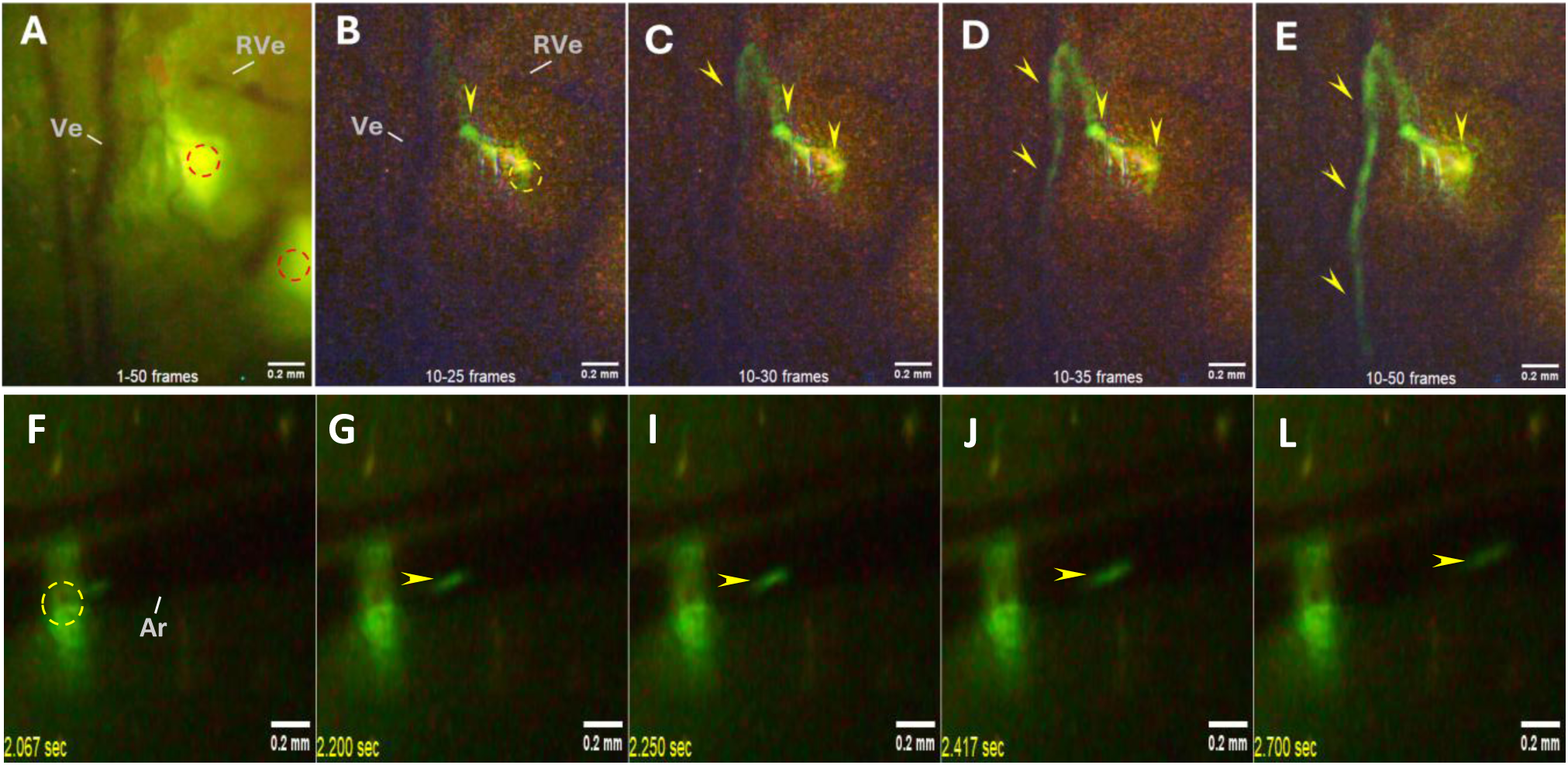
Intravascular detection of saliva released by *Lutzomyia longipalpis* during blood feeding. Upper panel: Visualization of a salivary jet emerging from the feeding site and penetrating an adjacent venule. (A) Representative image of the feeding site showing regions of saliva deposition on the skin surface and associated blood vessels. (B–E) Sequential frames illustrating the trajectory of labeled saliva from the dermal feeding site into the venular lumen. Lower panel: Intravascular transport of saliva within a venule (F–L). Sequential images show the displacement of fluorescently labeled saliva along the vessel, consistent with carriage by blood flow. Annotations: Saliva deposition (red circle); Mouthpart region (yellow circle); Venule (Ve); Ruptured venule (RVe); Saliva trajectory/tracking (yellow arrows).

## Discussion

### Mechanical and Microvascular Aspects of Blood Feeding in *Lu. longipalpis*

Relatively few studies have addressed the blood-feeding mechanisms of phlebotomine sand flies in detail. To date, the only comprehensive description encompassing the sequential phases of blood feeding on a live host was conducted nearly a century ago in the Old World species *Phlebotomus argentipes* by (36).

Since that seminal work, phlebotomine sand flies have been broadly classified as pool-feeding (telmophagous) arthropods, acquiring blood from dermal vessels through lacerations produced by their serrated mouthparts (36), (10). By employing new technologies in intravital microscopy and image analysis, the findings presented in this study provide valuable insights into the mechanical and physiological interactions between *Lu. longipalpis* and the microvasculature of the host. These findings not only complement but also expand the classical work by (36) on *P. argentipes*, revealing both conserved and novel aspects of sand fly blood-feeding strategies.

A particularly notable behavioral pattern observed during the initial phase of feeding was the occurrence of “scissor-like” movements of the labrum and hypopharynx during mouthpart penetration (Figs2A and B; S1Video). These movements are likely to contribute to dermal tissue damage and vascular laceration, facilitating blood extravasation. While Shortt and Swaminath did not describe such motions, they concluded that blood was rarely drawn directly from intact vessels; rather, they suggested that capillary damage caused by the proboscis tip resulted in the pooling of blood in the dermis, which was subsequently ingested. The formation of hemorrhagic pools and occasional blood splashes recorded during our experiments suggests that tissue disruption is an integral part of this process; however, passive blood leakage alone is insufficient to sustain feeding. Instead, successful blood uptake depends on precise mouthpart coordination and repeated adjustments at the feeding site.

Feeding in *Lu. longipalpis* proceeded through two distinct phases: an erratic probing phase, often marked by unsuccessful or inconsistent blood intake despite visible hemorrhage, followed by a stable engorgement phase. The transition between these phases was frequently associated with the recruitment or opening of one or more small dermal vessels, referred to here as “feeder vessels” that would deliver the blood directly to the insect’s mouthparts (Figs 2E and F; G and H; S2Video).

In some instances, engorgement began immediately after the flow from these vessels was established. These observations suggest that successful feeding requires not only mechanical disruption but also a favorable microvascular response by the host.

In contrast, (36) noted a delay between the insertion of the mouthparts and the onset of blood intake. Although they did not provide a vascular explanation for this lag, our findings suggest that it may reflect an initial vasoconstriction triggered by mechanical trauma, followed by vascular relaxation or rupture that enables sustained blood flow. This sequence of microvascular responses - vasoconstriction followed by targeted vasodilation or vessel disruption - has not been previously described and may be critical for feeding success in sand flies (Fig 3).

The “feeder vessel” observed in our study appeared to be ruptured but not collapsed, exhibiting intermittent blood flow toward the insect’s mouthparts. This condition likely ensures a steady and localized blood supply during the engorgement phase and may reflect a specific vascular response elicited by sand fly probing behavior. Such dynamic interactions were not previously addressed in the literature, which had focused primarily on gut distension and the mechanical aspects of ingestion.

### Salivary Secretion pattern during Blood Feeding

Acridine Orange (AO) proved to be an effective vital dye for labeling the saliva of *Lu. longipalpis*, enabling detailed visualization of salivary activity throughout feeding. Similar applications of AO have been previously reported in hematophagous insects such as triatomines and bed bugs (37),(38). In our experiments, all *Lu. longipalpis* individuals fed on AO-supplemented sucrose displayed strong labeling of their salivary glands. In some dissected specimens, a reduction in gland size and fluorescence intensity was observed post-blood feeding (Figs 4C and D), indicating reduction of salivary contents. This finding is consistent with previous studies, which show a ∼95% reduction in total salivary gland proteins in *Lu. longipalpis* after blood feeding on hamsters (39).

AO is a weak base that crosses cell membranes under physiological pH conditions (e.g., blood and hemolymph, ∼7.2–7.4) and accumulates in acidic compartments (40),(41). It is an effective accumulation in the salivary glands of *Lu. longipalpis,* as in *Rhodnius prolixus* and *Cimex lectularius* supports the hypothesis that these glands have an acidic environment (37)(38). In our study, pH indicators (bromothymol blue and bromocresol purple) confirmed that *Lu. longipalpis* salivary content of has a pH close to 6.0 (Figs 4E and F).

The extensive release of saliva during the initial feeding stages was observed in *Lu. Longipalpis* mirrors patterns reported in other blood-feeding insects, such as triatomines and bed bugs (37) (38). This early salivation likely counteracts host responses triggered by tissue injury, including vasoconstriction and platelet aggregation, which could impede blood flow. The presence of vasodilators and anti-inflammatory molecules in sand fly saliva supports this hypothesis (42), (15). Since the initial feeding phase is also the most perceptible by the host (43), rapid salivation may help establish anti-nociceptive effects. In *Lu. longipalpis*, this could be mediated by salivary adenosine deaminase (44) or sodium channel-blocking activity, as described in triatomine saliva (45). Early suppression of host detection would reduce grooming behavior and the risk of feeding interruption or insect injury.

We also observed saliva being released through the opening of the salivary canal near the tip of the hypopharynx, particularly when the mouthparts crossed the superficial dermis. At these moments, a rapid change in saliva consistency was seen, with the secretion assuming a gelatinous appearance upon exposure to air (S6Video). This “gel-like saliva” phenomenon was observed in both AO-labeled and unlabeled insects, suggesting it is not an artifact of staining. Similar transitions from liquid to gel saliva have been described in phytophagous insects in response to oxygen exposure (46), (47) often involving oxidative polymerization of sulfhydryl-containing proteins (48). In these insects, gel saliva contributes to forming salivary sheaths that protect feeding stylets and minimize sap loss (49). Although we observed comparable changes, oxygen-dependent gelification in hematophagous insects has not been reported and warrants further investigation.

Post-feeding monitoring revealed that AO-derived fluorescence remained visible in the host skin for over an hour, consistent with the telmophagic feeding strategy of *Lu. longipalpis*, involving substantial dermal tissue disruption and the action of salivary enzymes such as hyaluronidase and nucleases. In contrast, vessel-feeding insects such as *R. prolixus* and *C. lectularius* exhibit much shorter fluorescence retention times at the feeding site, likely due to reduced tissue damage and the transport of saliva into blood vessels by circulation (33), (38).

In numerous experiments, the intensity and persistence of fluorescence at the feeding site hindered real-time visualization of salivation dynamics. To overcome this limitation, we applied the Z-Project or frame-by-frame subtraction in Fiji / ImageJ, which enables enhanced temporal resolution of salivary events through projection-based image processing. This analytical approach permitted the identification of two distinct phases of salivation: (1) an initial high-frequency phase occurring immediately after skin penetration, frequently associated with mouthpart repositioning during probing; and (2) a subsequent prolonged low-frequency phase, characterized by more rhythmic salivary release, averaging approximately one salivation peak every 3.6 s (Fig 5B). Further analysis revealed that each salivary ejection was preceded by a fluorescent halo surrounding the mouthparts - most prominently the labrum and hypopharynx - and was often followed by a rapid closure of the mouthparts. As feeding progressed, the trajectories of the salivary ejections penetrated progressively deeper into the dermis, likely reflecting the activity of salivary hyaluronidase in degrading components of the extracellular matrix, particularly hyaluronic acid.

These observations support the established view that sand fly saliva contains an array of bioactive molecules essential not only for blood feeding but also for *Leishmania* transmission. Hyaluronidase, found in *Lu. longipalpis*, *P. papatasi*, and other species facilitate the spread of saliva and parasites in host tissues (28), (6). Maxadilan, a potent vasodilator unique to *Lu. longipalpis*, acts on PAC1 receptors to increase blood flow and modulate host immunity (50), (51), (52). Salivary nucleases degrade extracellular DNA, including neutrophil extracellular traps (NETs), aiding parasite evasion and dissemination (53).

Additional components such as apyrases, lufaxin, yellow-related proteins, and D7-related proteins further disrupt host hemostatic and inflammatory pathways (15),(54),(55),(56). The synergistic action of these molecules enhances feeding success while promoting the survival of *Leishmania*. Notably, the observation of fluorescent saliva jets penetrating intact vessel walls without visible or hemorrhage aligns with findings in *R. prolixus*, where increased vascular permeability was demonstrated via intravital microscopy and plasma markers (33).These results suggest that sand fly saliva may cross vascular barriers through chemically mediated mechanisms involving vasoactive molecules such as nitric oxide rather than mechanical damage alone.

Although both *Lutzomyia longipalpis* and the argasid tick *Ornithodoros rostratus* are classically categorized as telmophagous arthropods, the feeding dynamics described for these taxa revealed functional principles alongside marked mechanistic divergences. In both species, blood acquisition relies on disruption of the dermal microvasculature and on the establishment of an extravascular hemorrhagic microenvironment that must be actively maintained through sustained salivary activity.

In *O. rostratus*, feeding follows a highly stereotyped sequence of events in which the initial phase of hemorrhagic pool formation represents only ∼5% of the total host-contact period and is dominated by intense cheliceral activity coupled with abundant salivary secretion prior to the effective onset of blood uptake (57). By contrast, our observations in *L. longipalpis* indicate a more prolonged and dynamically modulated salivary deployment during probing and feeding. Collectively, these distinctions underscore that telmophagy does not constitute a uniform feeding strategy but rather encompasses a spectrum of adaptive solutions shaped by lineage-specific evolutionary trajectories within hematophagous arthropods.

## Conclusion

Our study refines the classical view of sand fly telmophagy by establishing that successful blood feeding in *L. longipalpis* is a coordinated process that integrates mouthpart mechanics, salivary release, and host microvascular modulation. Rather than arising from passive blood pooling alone, feeding relies on the targeted recruitment of dermal “feeder vessels” and salivation that dynamically reshapes the feeding microenvironment. These findings provide new insights into how feeding behavior enhances parasite transmission and highlight salivary factors as key targets for transmission-blocking strategies.

## Acknowledgments

The authors would like to acknowledge Mr. Cesar Nonato de Oliveira for his technical oversight of the sand flies colony

## Supporting information Captions

**Video S1-** Scissor-like mouthpart movements and blood splashes during probing by *Lutzomyia longipalpis*. Following skin penetration, the labrum and hypopharynx perform repeated penetration and withdrawal movements, remaining closely apposed but occasionally separating in characteristic scissor-like motions. During the initial probing phase, small hemorrhagic spots become visible on the skin surface, some of which are associated with active blood uptake (1.483 sec). At specific moments, these mouthpart structures close forcefully, generating small blood splashes that disperse from the mouthpart region (1.000 sec). The primary view of the video shows a frame that has been extracted to illustrate the detailed features of mouth parts. The Video is displayed at 30 fps to facilitate detailed observation. Arrows indicate: Labrum (La) and Hypopharynx (Hy).

**Video S2 -** Characteristic positioning of mouthparts during blood ingestion by *Lutzomyia longipalpis.* The opening view of the video consists of a frame selected to showcase the detailed features noted throughout this recording. The video shows the moments of closure (1.550 sec) and opening of the “feeding vessel (2.017 sec) during active blood uptake. Abbreviations: Labrum (La); Feeding vessel (Fv); Functional mouth (Fm); Hypopharynx (Hy). The circle in yellow highlights the functional mouth area.

**Video S3 -** Saliva emission and mouthpart movements during blood feeding by *Lutzomyia longipalpis* labeled with acridine orange (0.1%). The video documents salivary release and associated mouthpart dynamics during blood feeding on the ear skin of a hairless mouse. The Video is displayed at 30 fps to facilitate detailed observation and the opening view of the video features a frame that has been taken to highlight the mouthpart region (Red arrow) and sites of saliva emission (white arrows).

**Video S4-** Jet-like saliva emission at the feeding site of *Lutzomyia longipalpis*. The video shows a directed jet of saliva emerging from the feeding site in close proximity to the wall of an adjacent blood vessel within the host tissue. The first frame shows a reversed image of the insect feeding site generated using the “Invert command” in ImageJ. The arrows point to the Saliva jets (Sa), Saliva accumulation in the tissues (*) and the host venule used during feeding (Ve). The subsequent frames correspond to post-processed image sequences derived from a 2,300-frame video, obtained through application of the Grouped Z Project (SUM projection) at intervals of every 10 frames. The Video is displayed at 30 fps to facilitate detailed observation.

**Video S5 -** Sandfly saliva observed inside blood vessels. Sequential images showing the release of fluorescently labeled saliva along the vessel following the blood flow. The first frame shows a reversed image of the insect feeding site generated using the “Invert command” in ImageJ. The Video is displayed at 30 fps to facilitate detailed observation and Arrows indicate: Venule (Ve); The Arteriole (Ar) in the feeding area is surrounded by saliva accumulated in the tissues (St) and the saliva (Sa), which is carried by the flow of blood.

**Video S6 -**The labeled saliva is released after the sandfly *Lutzomyia longipalpis* overpasses the host skin. The first frame of the video illustrates the details of the Saliva accumulation in the tissues (St) and saliva (Sa) being released in the vicinity of the area penetrated by the insect.

